# Clustering of CD3ζ is sufficient to initiate T cell receptor signaling

**DOI:** 10.1101/2020.02.17.953463

**Authors:** Yuanqing Ma, Yean J Lim, Aleš Benda, Jesse Goyette, Katharina Gaus

## Abstract

T cell activation is initiated when ligand binding to the T cell receptor (TCR) triggers intracellular phosphorylation of the TCR-CD3 complex. However, it remains unknown how biophysical properties of TCR engagement result in biochemical phosphorylation events. Here, we constructed an optogenetic tool that induces spatial clustering of CD3ζ chains in a light controlled manner. We showed that spatial clustering of the CD3ζ intracellular tail alone was sufficient to initialize T cell triggering including phosphorylation of CD3ζ, Zap70, PLCγ, ERK and initiated Ca^2+^ flux. In reconstituted COS-7 cells, only Lck expression was required to initiate CD3ζ phosphorylation upon CD3ζ clustering, which leads to the recruitment of tandem SH2 domain of Zap70 from cell cytosol to the newly formed CD3ζ clusters at the plasma membrane. Taken together, our data suggest that clustering of the TCR can initialize proximal TCR signaling and thus constitute a biophysical mechanism of TCR triggering.

## Introduction

T cell activation is initialized by peptide-bound major histocompatibility complex (pMHC) molecules engaging the αβ chain of the TCR. The TCR is constitutively associated with CD3 dimers – CD3εγ, CD3εδ, CD3ζζ – and upon ligation of the αβTCR, the immunoreceptor tyrosine-based activation motifs (ITAM) of CD3 molecules are phosphorylated by the tyrosine kinase Lck. TCR-CD3 phosphorylation is regarded as the first biochemically detectable signal of T cell activation and is termed TCR triggering. The exact mechanisms of how ligand binding to the extracellular domain of the αβTCR triggers CD3 phosphorylation of the intracellular motif remains unclear. Ligand binding may induce conformational changes (Gil et al., 2002, Lee et al., 2015) to facilitate coordinated re-arrangements of the TCR-CD3 subunits. However, conformational changes cannot explain TCR signaling induced by antibodies or triggering of simpler versions of the TCR such as chimeric antigen receptors (CARs). For example, T cells that express the extracellular and transmembrane domains of the co-receptor CD8 fused to the intracellular domain of CD3ζ respond to ligation in a similar manner as the TCR-CD3 complex, including activation of the canonical TCR signaling pathways (Irving and Weiss, 1991). The ability of CARs to elicit T cell responses analogous to the native TCR by integrating into the T cell signaling network has led to the clinical success of CAR T cell therapies (Kalos et al., 2011). It also raises the fundamental question of whether other mechanisms apart from conformational changes are involved in TCR triggering.

Emerging evidence suggests that TCR clustering is required and/or regulates TCR triggering. First, while both monomeric and oligomeric pMHC bind to the TCR, only oligomeric ligands trigger TCR signaling, suggest that ligand-induced receptor clustering is required for activation (Cochran et al., 2000, Boniface et al., 1998). Other studies also suggested that antibody-induced receptor crosslinking was responsible for CAR triggering (Irving and Weiss, 1991, Eshhar et al., 1993, Letourneur and Klausner, 1991). Second, it is well-documented that the TCR forms nano- and micro-clusters immediately after engagement with pMHC molecules or activating antibodies on cell or engineered surfaces (Yokosuka et al., 2005, Ike et al., 2003, Varma et al., 2006). In activated T cells, proximal signaling molecules such as Zap70, LAT, SLP-76 and PLCγ associate with TCR clusters (Yokosuka et al., 2005, Varma et al., 2006). A DNA hybridization-based CAR revealed that a single ligand receptor interaction can induce receptor clustering and initiate signaling (Taylor et al., 2017). Third, we found that only dense TCR clusters had a high signaling efficiency, suggesting a direct relationship between receptor clustering to signaling (Pageon et al., 2016).

To directly test whether clustering alone can induce receptor triggering, it is advantageous to bypass the engagement of external oligomeric ligands and directly induce receptor clustering. A previous study (Spencer et al., 1993) employed a chemically induced oligomerization system to demonstrate that the induced clustering of membrane-anchored CD3ζ chain was sufficient to trigger the activation of NF-AT, Oct/OAP and AP-1 transcription factors. In their system, CD3ζ oligomerization was initialized by the membrane-permeable molecule FK1012 that induces homo-dimerization of FK506-binding protein (FKBP) in the cell. By comparing the activation of CD3ζ linked to one, two or three copies of FKBP, the authors concluded that valency of the receptor binding to FK1012 and the consequent oligomerization state was synergistic to receptor activation.

To control clustering with higher flexibility, we developed an optogenetic CD3ζ construct that self-associates non-invasively upon irradiation with blue light and without the use of ligands. While optogenetic techniques are well-established in neuroscience, these tools have recently been applied to T cell biology. For instance, two studies have employed optogenetics to control the ligand-receptor binding lifetime to test the kinetic proof reading model of T cell activation (Tischer and Weiner, 2019, Yousefi et al., 2019). In the current study, we employed optogenetics to control the clustering of CD3ζ by tagging the photoreceptor cryptochrome 2 (Cry2) to the C terminus of CD3ζ. This construct forms homo-oligomers when the Cry2 chromophore is excited by blue light (Bugaj et al., 2013). Using this tool, we demonstrated that the spatial clustering of the intracellular domain of the CD3ζ chain was sufficient to trigger downstream T cell activation signaling events. We reconstitute light-induced CD3ζ triggering in non-T cells to demonstrate that clustering *per se* constitutes a biophysical mechanism of signal initiation. We propose that in T cells, multiple TCR triggering mechanisms can co-exist, enabling T cells to probe antigens under a large number of conditions.

## Results

### An optogenetic tool to control the clustering of CD3ζ

We aimed to address whether clustering of the intracellular domain of CD3ζ is a regulatory element in TCR triggering. To avoid any association of CD3ζ with endogenous TCR subunits, we replaced the extracellular and transmembrane domain of CD3ζ with the membrane anchor of Lck so that the intracellular domain of CD3ζ is attached to the plasma membrane via palmitoylation and myristoylation groups (Zlatkine et al., 1997). Previous studies showed that proteins with this membrane anchor diffuse as monomers in the plasma membrane (Douglass and Vale, 2005, Triffo et al., 2012). To control the clustering of the CD3ζ non-invasively and without ligand binding, we linked the light sensitive photolyase homology region (PHR) domain (1-498 AA) of Cry2 (Bugaj et al., 2013) to the C-terminus of CD3ζ (**Fig. 1a**). Under blue light illumination, Cry2 self-associates into clusters in a reversible manner (Duan et al., 2017). We fused the construct to mCherry to visualize the protein under 594 nm excitation and named the final construct ‘light-induced clustering of CD3ζ’ (LIC-CD3z, **Fig. 1a**). As a control, we made the equivalent construct but without the Cry2 sequence (**Fig. 1b**, named LIC-CD3z-delCry2). Both constructs expressed well in Jurkat or COS-7 cells and were mainly targeted to the plasma membrane, as expected (data not shown). Confocal imaging confirmed that LIC-CD3z clustered immediately after irradiation with blue light while LIC-CD3z delCry2 did not (**Fig. 1c**).

**Figure 1.**
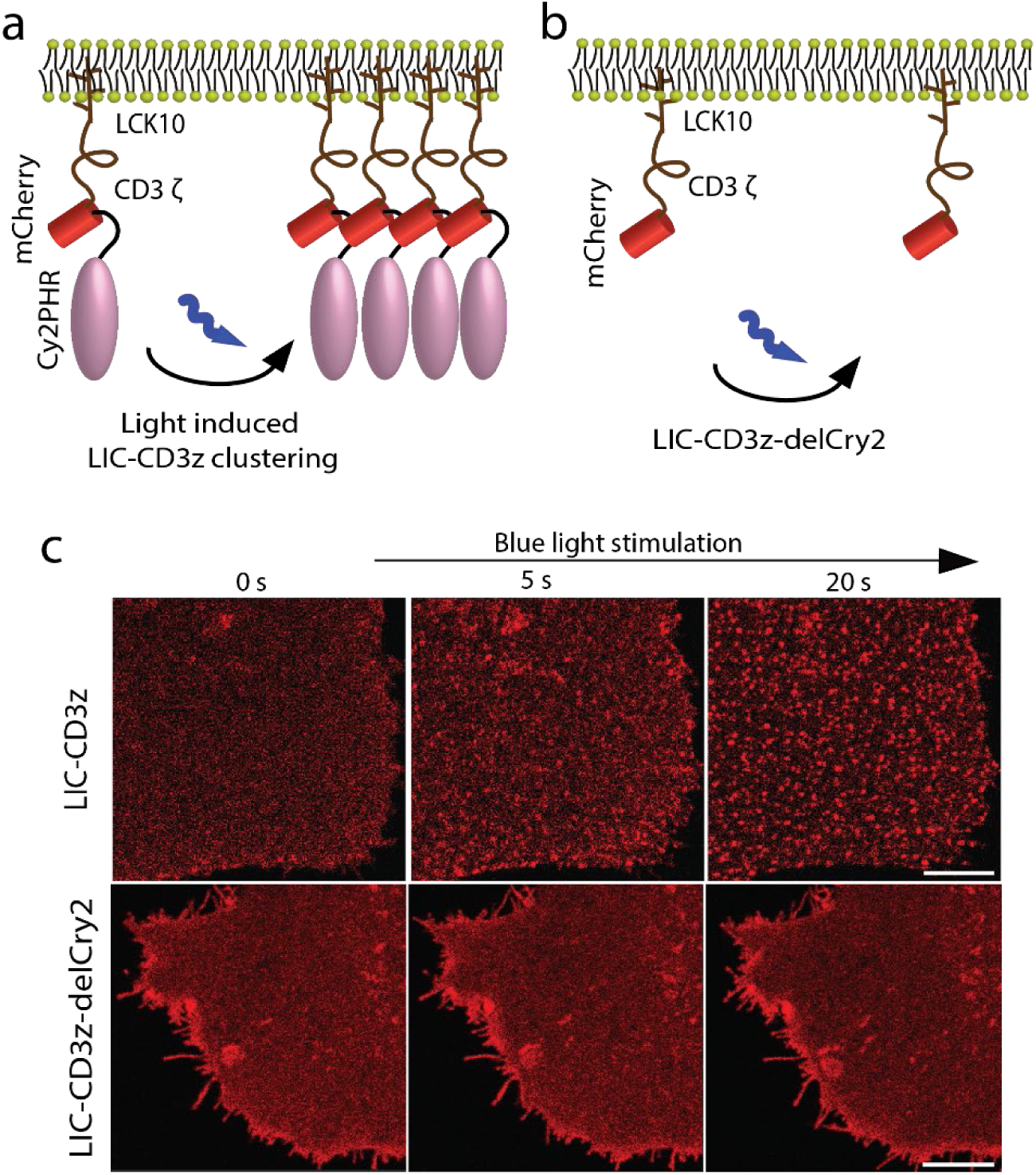
Light-induced clustering of CD3ζ. **a**. Schematic of the light-induced construct termed LIC-CD3z consisting of a membrane anchor (first 10 amino acids of Lck), the cytosolic tail of CD3ζ, mCherry and the light sensitive domain PHR of Cry2. Prior to illumination, LIC-CD3z diffuses freely in the plasma membrane as monomers. Upon illumination with blue light, LIC-CD3z self-associates into clusters. **b**. Schematic of the control construct, termed LIC-CD3z-delCry2, which lacks the Cry2 domain and is light insensitive. Schematics (a-b) are not to scale. **c**. Confocal images of LIC-CD3z (top row) and in LIC-CD3z-delCry2 (bottom row) in COS-7 cells. Cells were irradiated with 488 nm light at low intensity and imaged by exciting mCherry with 594 nm light. Images are representatives of n = 5 experiments. Scale bar =3µm.

### Clustered LIC-CD3z induces Ca^2+^ flux independent of the TCR complex

To verify if LIC-CD3z was signaling competent, we first investigated whether CD3ζ clustering was sufficient to trigger downstream signaling events, measured here as Ca^2+^ fluxes. We transfected LIC-CD3z into αβTCR-deficient T cells, Jurkat 76 cells, that have essentially no endogenous CD3 expression on the cell surface (Heemskerk et al., 2003, Knies et al., 2016). Thus, any signaling exhibited in these cells would be restricted to LIC-CD3z and would not involve other components of the TCR complex. A genetically encoded Ca^2+^ sensor, G-GECO (Zhao et al., 2011), was co-transfected as a readout of T cell activation. Here, the 488 nm laser excited G-GECO and activated Cry2, such that clustering of LIC-CD3z and time-lapse imaging of G-GECO was performed simultaneously. To confirm that any signaling was initialized by CD3ζ clustering, two control constructs were tested under identical conditions (**Fig. 2a**): LIC-CD3z-delCry2, which lacks the Cry2 domain (**Fig. 1b**) and is light insensitive (**Fig. 1c**) and LIC-CD3z Y-L, which has all six tyrosine residues in the three ITAMs of the CD3ζ chain replaced by leucine residues rendering it effectively a CD3ζ signaling-defective mutant. Time-lapse images (**Fig. 2b**) and movies (**Video 1**) showed that the clustering of LIC-CD3z caused Ca^2+^ influx in transfected Jurkat 76 cells ∼80 s into irradiation with blue light (**Fig. 2c**). In contrast, Jurkat cells expressing LIC-CD3z-delCry2 or LIC-CD3z Y-L exhibited no measurable Ca^2+^ fluxes, suggesting that the observed Ca^2+^ signaling was triggered by CD3ζ clustering and required phosphorylated ITAMs.

**Figure 2.**
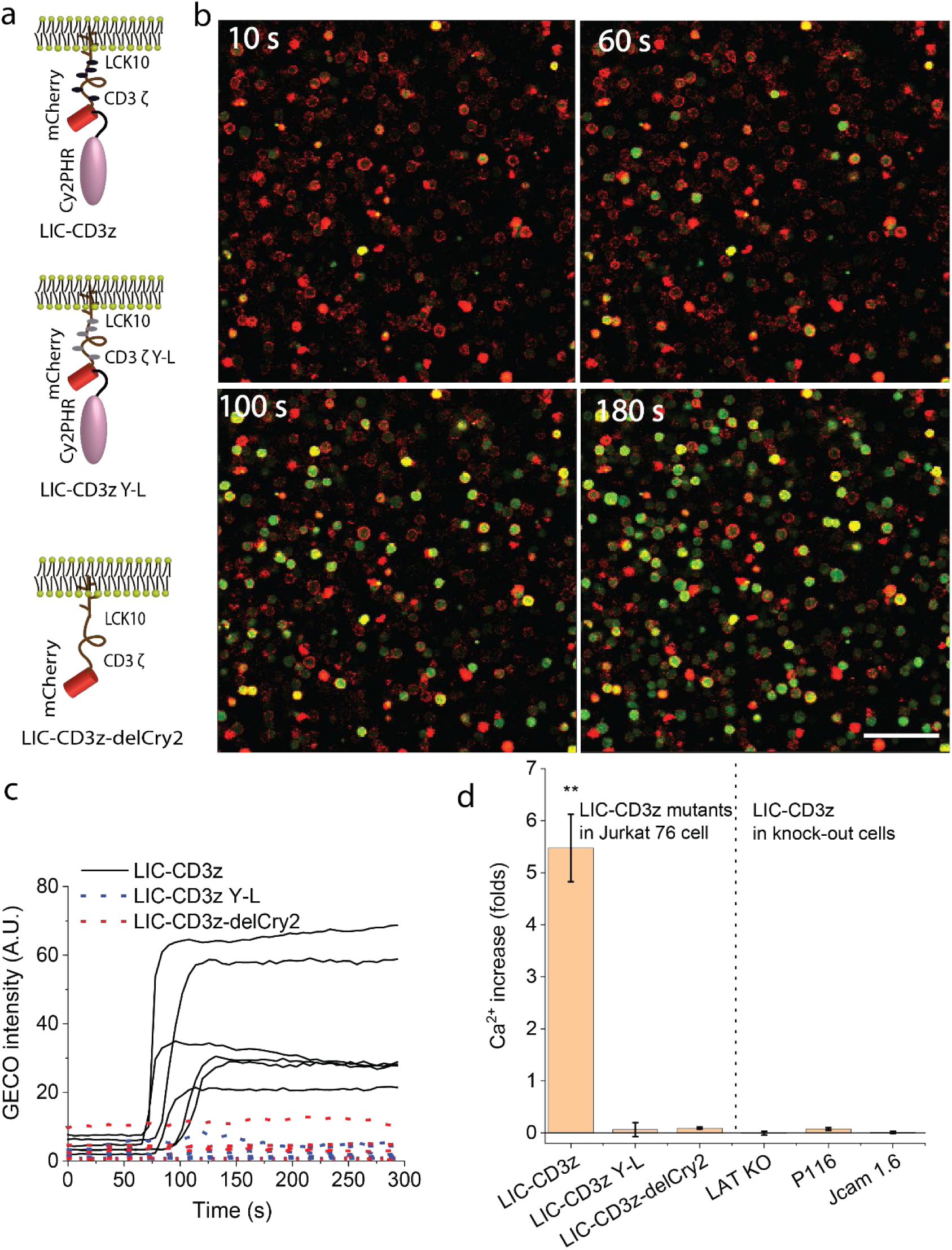
LIC-CD3z clustering induces Ca^2+^ influx in Jurkat cells. **a**. Schematics of LIC-CD3z (top), signaling incompetent LIC-CD3z Y-L(middle), and light insensitive LIC-CD3z-delCRY2 (bottom). **b**. Confocal images of Ca^2+^ influx in Jurkat 76 cells co-transfected with LIC-CD3z (red) and Ca^2+^ sensor G-GECO (green). Images were taken at the indicated time points after irradiation with blue light. **c**. G-GECO intensity traces over time for single cells expressing LIC-CD3z (solid line), LIC-CD3z-delCRY2 (red dotted line) and LIC-CD3z Y-L (blue dotted line). Scale bar = 150 µm. **d**. Quantification of Ca^2+^ influx, as fold increase over baseline level, in Jurkat 76 cells expressing LIC-CD3z, LIC-CD3z Y-L and LIC-CD3z-delCRY2, and LIC-CD3z expressed in Jurkat cells deficient of LAT (LAT KO), ZAP70 (P116) or Lck (JCam 1.6). In d, data are mean and standard error of n = 30 cells. ** p<0.01 (one-way ANOVA with Bonferroni post hoc test).

The canonical signaling pathway of TCR triggering follows a sequence of events that begins with the phosphorylation of ITAMs by Lck, followed by membrane recruitment of Zap70 to the phosphorylated ITAMs, where Zap70 becomes activated by both transphosphorylation (Chan et al., 1995) and phosphorylation by Lck, and the recruitment and tyrosine phosphorylation of LAT. We therefore enquired whether LIC-CD3z clustering engages the same signaling pathway. For this we repeated the Ca^2+^ flux experiment in Jurkat-derived cell lines lacking one of the proximal signaling molecules: JCam1.6 (Lck-deficient), P116 (Zap70-deficient), and a CRISPR/CAS9-gene edited LAT-knock out cell line. LIC-CD3z clustering did not induce Ca^2+^ flux in any of these cell lines (**Fig. 2d**), suggesting that LIC-CD3z clustering is likely to trigger the canonical TCR activation pathway. To confirm this, we performed Western blotting on LIC-CD3z-transfected Jurkat 76 cell lines to examine the phosphorylation of typical downstream signaling molecules. Cells were irradiated for 45 seconds and kept in the dark for 1-5 min to prevent continuous LIC-CD3z clustering prior to cell lysis. We found that CD3ζ (at Y142), Zap70 (at Y319) and phospholipase C-γ1 (PLCγ, at Y783) were phosphorylated within the first minute after light exposure, and the extracellular signal regulated kinase (ERK1/2) after ∼5 min (**Fig. 3**). Activated PLCγ hydrolyses PIP_2_ to diacylglycerol (DAG) and inositol 1,4,5 trisphosphate (IP_3_), which releases Ca^2+^ from the endoplasmic reticulum and induces further flux through membrane Ca^2+^ channels (Lewis, 2001). It is thus likely that the observed Ca^2+^ flux was caused by PLCγ activation. ERK1/2 phosphorylation is required for the activation of T cell effector function such as interleukin-2 (IL-2) secretion (Whitehurst and Geppert, 1996). Taken together, the data suggest that clustering of the cytosolic tails of CD3ζ in the plasma membrane of T cells is sufficient to initiate early TCR signaling in a similar manner as pMHC-TCR ligation. Since the CD3ζ tails are unstructured, it is unlikely that conformational changes are mechanistically involved in signal initiation with LIC-CD3z. The need for the ITAM domains in LIC-CD3z and the lack of Ca^2+^ fluxes in Lck-deficient cells suggest that Lck is responsible for signal initiation. It is therefore possible that Lck phosphorylation of clustered substrates is more efficient than phosphorylation of monomeric substrates.

**Figure 3.**
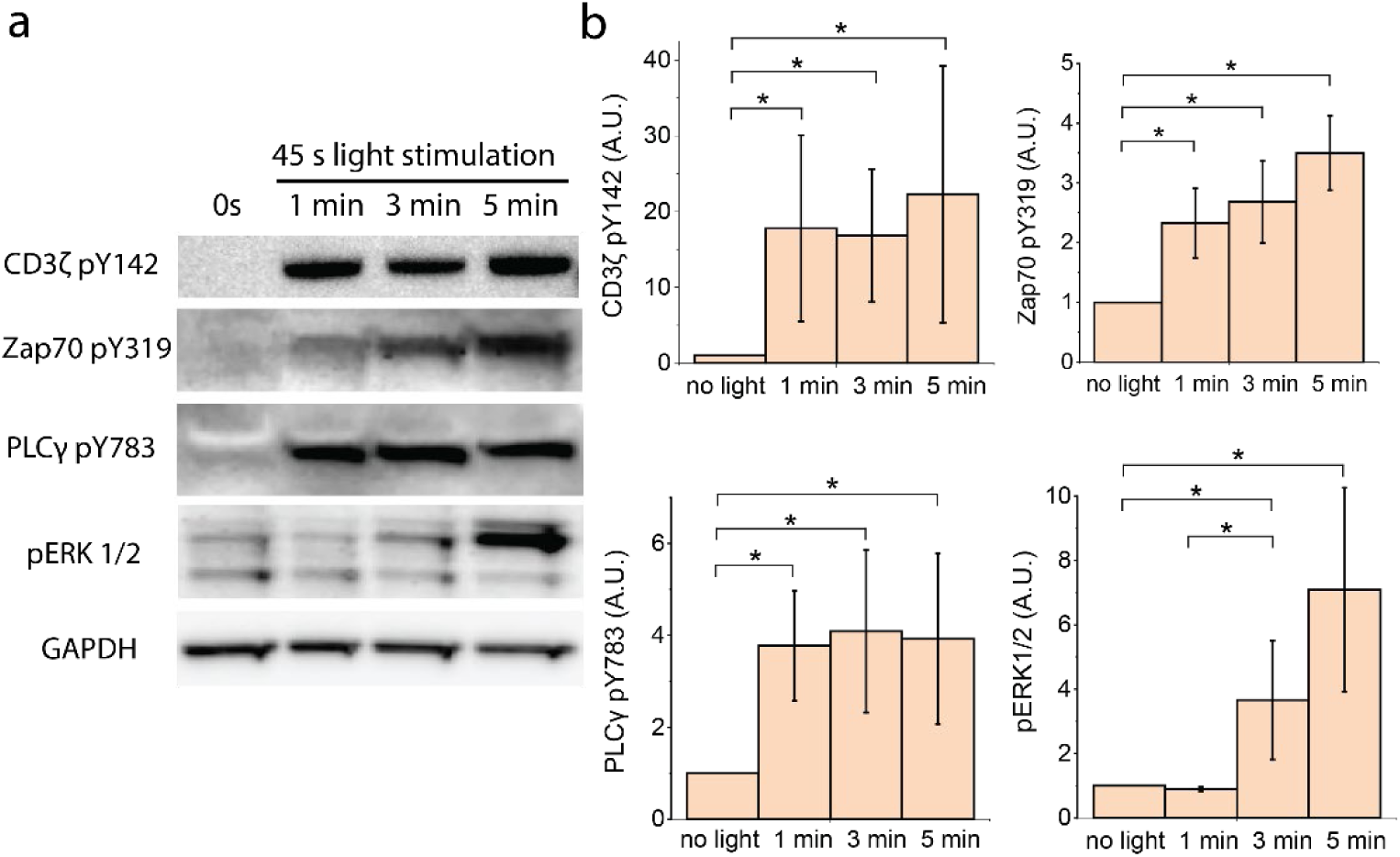
Western blot of TCR proximal signaling molecules in LIC-CD3z-transfected Jurkat 76 cell line. Western blot images (a) and quantification (b) of Jurkat 76 cell line transfected with LIC-CD3z and treated, or not, with blue light for 45 s followed by incubation in the dark for indicated time period (minutes). GAPDH was used as a loading control. Data are n=3 independent experiments, * p<0.05 (one-way ANOVA with Bonferroni post hoc test).

### Lck phosphorylated clustered LIC-CD3z efficiently in reconstituted COS-7 cells

To identify the minimal requirements for CD3ζ triggering and understand how clustering regulates CD3ζ phosphorylation, we reconstituted T cell signaling in a non-hematopoietic cell system. We co-transfected LIC-CD3z and wild-type Lck fused to green fluorescent protein (Lck GFP) into COS-7 cells and analyzed the LIC-CD3z ITAMs phosphorylation by immunostaining (**Fig. 4a**). Since the constructs were contained on two separate plasmids, transfected cells contained polyclonal populations with a range of expressions of each construct (**Supplementary Fig. 1**). For quantification, we therefore normalized phosphorylation levels to the expression levels of the CD3z construct (**Fig. 4b**). In cells that expressed LIC-CD3z but lacked Lck expression, LIC-CD3z was not phosphorylated (**Fig. 4a** red * labelled). In contrast, cells that expressed both constructs were stained by an Alexa647-tagged antibody against pY142 on ITAM3 (white cells in the merged image in **Fig. 4a**). However, we found no difference in ITAM3 phosphorylation levels before and after light-induced clustering of LIC-CD3z (**Fig. 4b** column 1 *versus* 2), suggesting that in the absence of key T cell phosphatases such as CD45 and SHP-1, Lck activity was sufficiently high to phosphorylate ITAMs regardless of the clustering state of CD3ζ. However, we found that light-treated LIC-CD3z resulted in significantly more phosphorylation than light-treated LIC-CD3z-delCry2 (**Fig. 4b** column 1 *vs* 3), ruling out that light treatment *per se* impacted on ITAM phosphorylation.

**Figure 4.**
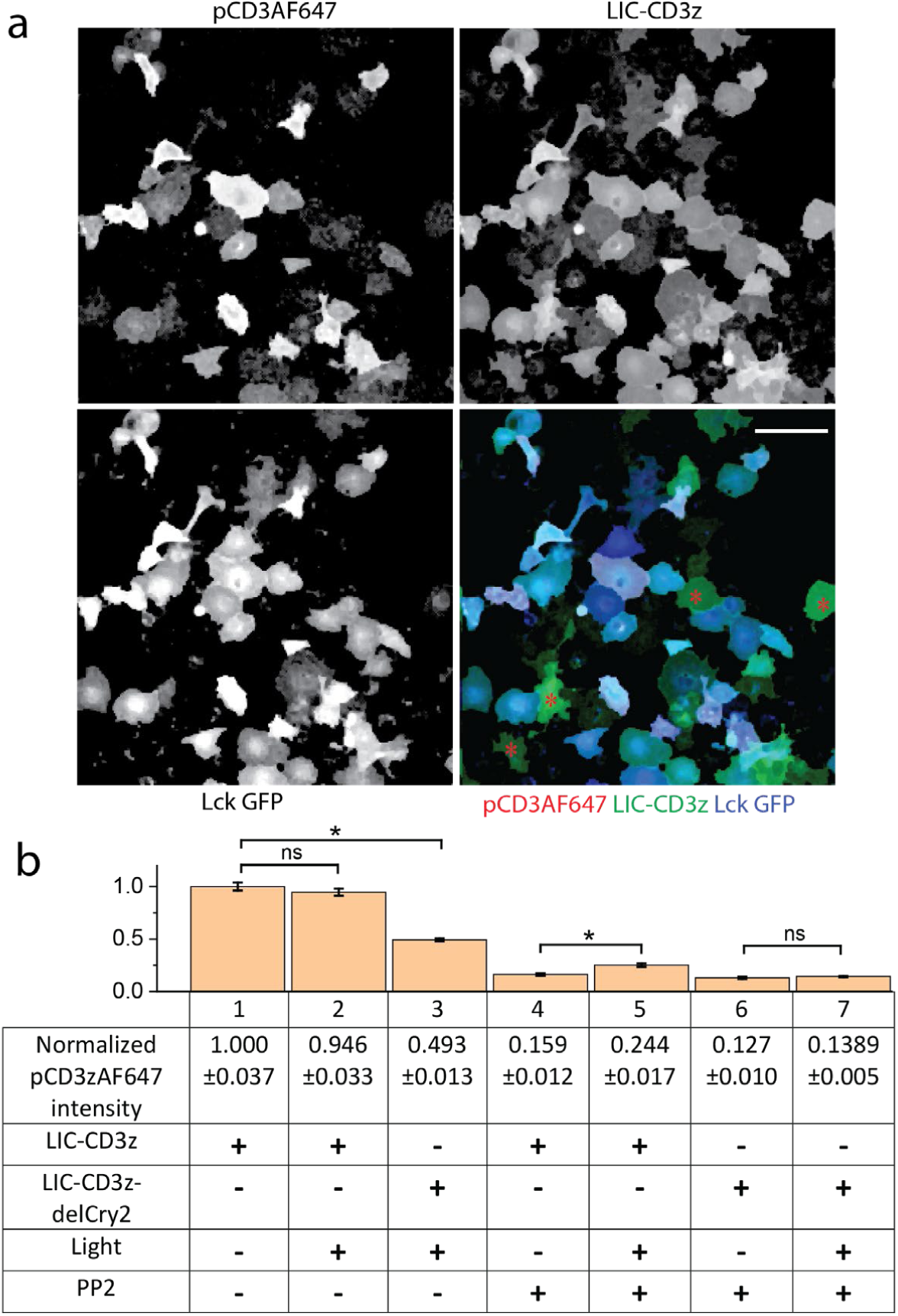
ITAM phosphorylation of LIC-CD3z and LIC-CD3z-delCry2 in COS-7 cells. a, Representative intensity images of individual and merged channels of COS-7 cells co-transfected with LIC-CD3z (green) and Lck-GFP (blue) and immune-stained with Alexa647-tagged antibodies recognizing Y142 in CD3ζ (red). Red asterisks (*) labelled cells that expressed only LIC-CD3z and exhibited no pCD3zAF647 staining. Scale bar = 150 µm. **b**. Bar graph and table of pCD3zAF647 intensity normalized to LIC-CD3z or LIC-CD3z-delCry2 expression levels (+/-) in cells that were treated or not treated with light (+/-) and were pre-treated or not treated with 25 µM of PP2 (+/-). Normalized pCD3zAF647 intensity in cells expressing LIC-CD3z without light or PP2 treatment were set to 1. * p<0.05, ns is not significant (one-way ANOVA with Bonferroni post hoc test).

To reduce basal phosphorylation, we incubated COS-7 cells overnight with 25 µM of the Lck inhibitor PP2, which was washed out 1 minute prior to light treatment. Our results showed that ITAM phosphorylation was substantially reduced compared to untreated cells (**Fig. 4b** column 1 *vs* 4). When LIC-CD3z clustering was induced with light in PP2-treated cells, the level of ITAM phosphorylation was increased significantly from 0.159 ± 0.012 to 0.244 ± 0.017 (column 4 and 5). In contrast, the non-clustering mutant LIC-CD3z-delCry2 displayed no significant change in phosphorylation under the same conditions (column 6 and 7). These experiments showed that firstly, when only LIC-CD3z was expressed in reconstituted COS-7 cells, there was a minimal level of phosphorylation by endogenous kinases. Secondly, when the kinase Lck was introduced, the basal kinase activity of Lck was sufficient to cause substantial LIC-CD3z phosphorylation. Thirdly, when Lck activity was controlled by PP2, clustering of LIC-CD3z facilitated increased phosphorylation by Lck. This suggests that under conditions where Lck activity is kept in check (either pharmacologically (**Fig. 4b**) or via the presence of endogenous phosphatase (**Fig. 2**), substrate clustering can be sufficient to shift the balance to a net increase in ITAM phosphorylation.

### LIC-CD3z clustering causes cytosol to plasma translocation of Zap70 in COS-7 cells

To measure CD3ζ triggering more directly, we used the tandem SH2 domain of Zap70 fused to mCherry (Zap70 tSH2) as a sensor for CD3ζ ITAM phosphorylation (Ottinger et al., 1998, Mukhopadhyay et al., 2013). In the absence of ITAM phosphorylation, Zap70 tSH2 diffuses freely in cell cytosol. Upon receptor trigging, the sensor binds to phosphorylated ITAMs and thus translocates to the plasma membrane (**Fig. 5a**). In agreement with the phosphorylation experiment, we noticed that in cells expressing higher levels of Lck, Zap70 tSH2 was already recruited to the cell plasma membrane prior to light activation of LIC-CD3z. Therefore, we conducted 3-colour confocal imaging of Zap70 tSH2, Lck GFP and LIC-CD3z fused to YFP in COS-7 cells with medium or low Lck expression level (**Fig. 5b**). By imaging the cross-section of cells, we could visualize and quantify the translocation of Zap70 tSH2 from the cytoplasm (center of cells) to the plasma membrane (edge of cells, **Fig. 5b** and **Video 2, 3**). To quantify Zap70 tSH2 association with the plasma membrane, we plotted the intensity ratio of regions representing the plasma membrane and the cytoplasm (**Fig. 5c**). While the association of the LIC-CD3z-YFP to the plasma membrane was constant (**Fig. 5d**), Zap70 tSH2 shifted from the cytoplasm to the plasma membrane upon CD3ζ clustering. The rapid association of Zap70 tSH2 with the plasma membrane after the start of imaging indicated that light-induced clustering enhanced ITAM phosphorylation above baseline levels.

**Figure 5.**
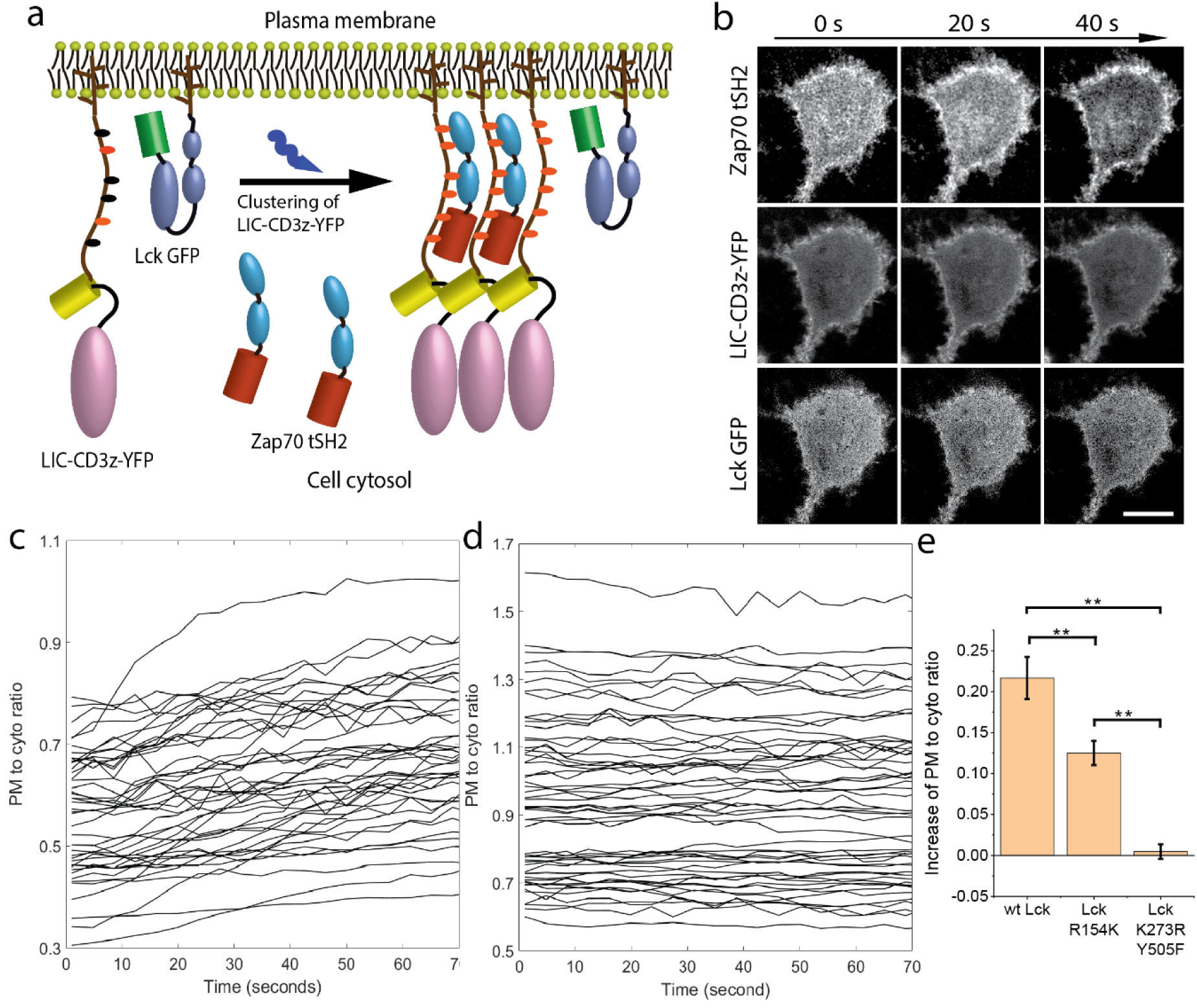
Reconstituting and imaging CD3ζ triggering in COS-7 cells. **a**. Schematics of reconstituting CD3ζ triggering in COS-7 by co-expressing LIC-CD3z-YFP, Lck GFP and the mCherry-tagged tandem SH2 domain of Zap70 (Zap70 tSH2). Prior to CD3ζ triggering, Zap70 tSH2 diffuses freely in the cytosol. Upon light-induced clustering, CD3ζ triggering is read-out as the translocation of Zap70 tSH2 from the cytosol to plasma membrane since Zap70 tSH2 binds specifically to phosphorylated ITAMs on the LIC-CD3z-YFP. **b**. Representative live cell confocal images of light-induced clustering of LIC-CD3z-YFP (top row), distribution of Zap70 tSH2 (middle) and Lck GFP (bottom). Scale bar = 20 µm. **c-d**. Quantification of Zap70 tSH2 (c) and LIC-CD3z-YFP (d) intensity at the plasma membrane relative to the cytosol immediately after irradiation. Each line represents one cell. **e**. Quantification of Zap70 tSH2 recruitment to the plasma membrane for cells expression wild-type Lck, Lck with a mutation in the SH2 domain (Lck R154R) or open and kinase dead Lck (Lck K273R Y505F). Zap70 tSH2 intensity ratio is shown as a fold-change relative to the first four imaging frames. Data are mean and standard error of n = 40 cells. ** p< 0.01 (one-way ANOVA with Bonferroni post hoc test).

To examine the role of Lck in Zap70 tSH2 recruitment to the membrane, we performed similar experiments using different Lck mutants with and without light-induced clustering of LIC-CD3z-YFP (**Fig. 5e**). The SH2 domain in Lck facilitates interactions with phospho-tyrosine residues, including phosphorylated ITAMs (Pellicena et al., 1998, Lewis et al., 1997). To determine the contribution of these interactions, we used a Lck version with a mutation in the SH2 domain (R154K Lck). We reasoned that Lck transiently interacting with CD3ζ *via* the SH2 domain may enhance phosphorylation efficiency in clusters as it increases local Lck concentration and residence time. Indeed, expressing R154K Lck resulted in less Zap70 tSH2 recruitment after CD3ζ clustering compared to cells expressing wild-type Lck (**Fig. 5e** and **Supplementary Fig. 1a, b**). This suggests that the SH2 domain of Lck was not mandatory but can enhance CD3ζ phosphorylation of clustered LIC-CD3z-YFP. Expressing an open but kinase-dead version of Lck, Lck K273R Y505F resulted in no increase in Zap70 tSH2 recruitment, indicating no change in CD3ζ phosphorylation levels (**Fig. 5e** and **Supplementary Fig. 1c, d**).

After Zap70 tSH2 is recruited to the plasma membrane, it should bind to the phosphorylated LIC-CD3z and thus rearranged into clusters. By focusing directly on the plasma membrane adjacent to the coverslip, we could indeed observe the gradual formation of Zap70 tSH2 clusters in the membrane (**Fig. 6a** and **Video 4**). From the 3-colour confocal images of the plasma membrane, we calculated the Pearson coefficient (Pike et al., 2017) for Zap70 tSH2 and LIC-CD3z-YFP co-localization (0.29 ± 0.03), which was substantially higher than the coefficient for Zap70 tSH2 and Lck co-localization (0.08 ± 0.02, **Fig. 6b**). The latter result supports the notion that Lck-CD3ζ interactions are highly transient, as previously suggested (Pageon et al., 2016). To exclude the possibility that the double SH2 domain of Zap70 tSH2 outcompetes the single SH2 domain of Lck to bind to phosphorylated ITAM, we also performed Pearson coefficient analysis in cells expressing LIC-CD3z and Lck GFP without Zap70 tSH2, which displayed a lack of colocalization (**Fig. 6b** lane 3). We followed the formation of LIC-CD3z-YFP clusters with live cell imaging and noticed that LIC-CD3z-YFP clusters were formed first, followed by the recruitment of Zap70 SH2 to LIC-CD3z-YFP clusters (**Fig. 6c**), resulting in an increases in the Pearson coefficient over time (**Fig. 6d**).

**Figure 6.**
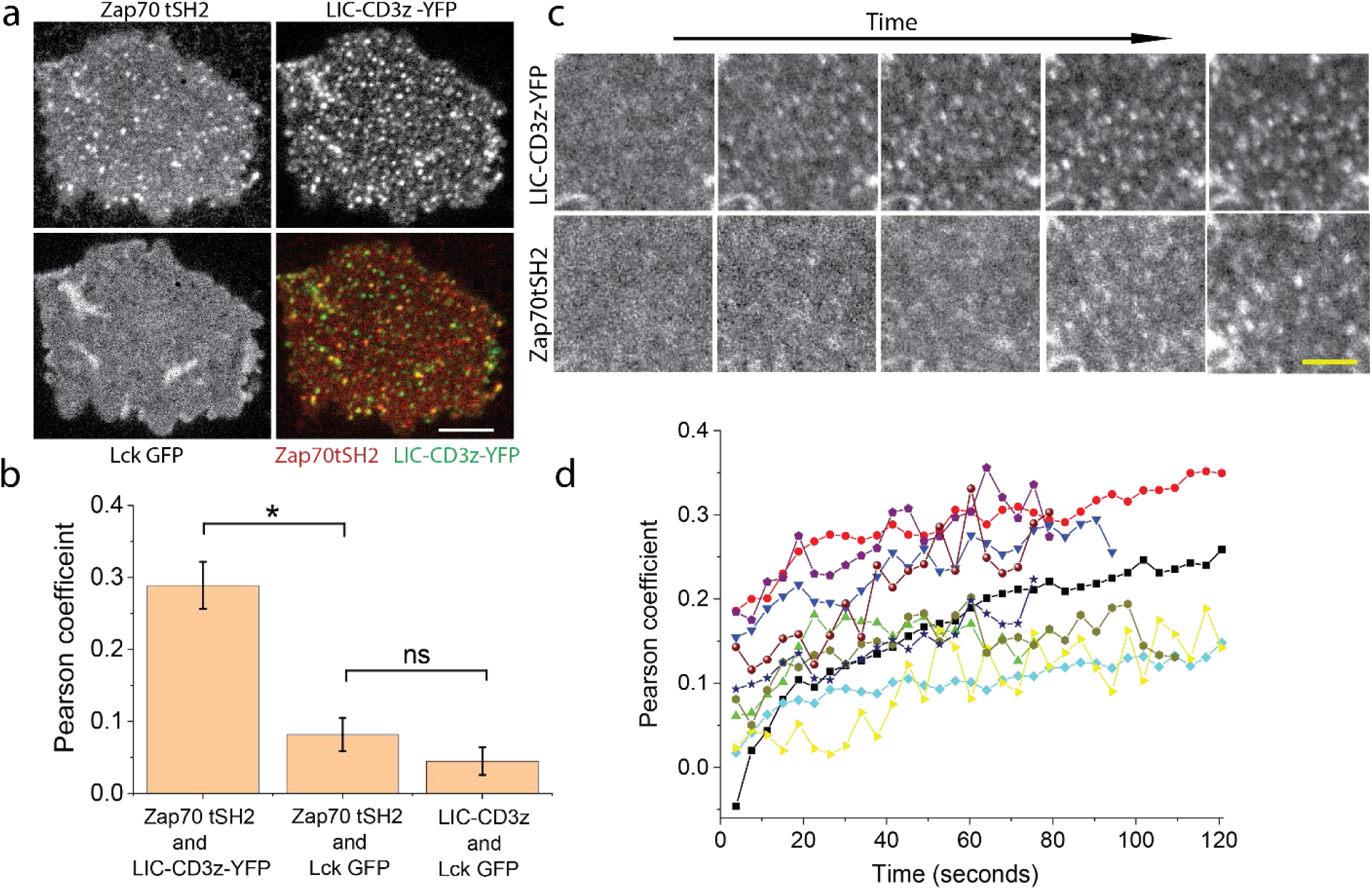
Zap70 tSH2 is recruited to LIC-CD3z-YFP clusters in reconstituted COS-7. COS-7 cells were reconstituted to co-express LIC-CD3z-YFP, mCherry-tagged Zap70 tSH2 and wild-type Lck GFP. **a-b**. Representative images of Zap70 tSH2, LIC-CD3z-YFP and Lck GFP colocalization in COS-7 cells immediately fixed after 2 minutes irradiation with blue light (a) and corresponding Pearson coefficient analysis (b). In b, the third column is Pearson coefficient values between LIC-CD3z and Lck GFP channel in COS-7 cells expressing only these two constructs. Scale bar = 10 µm. Data are mean and standard error of n = 20 cells. * p< 0.05, (one-way ANOVA with Bonferroni post hoc test). **c-d**. Representative images of the dynamic recruitment of Zap70 tSH2 to newly formed LIC-CD3z-YFP clusters in live COS-7 cells (c) and frame-by-frame Pearson coefficient values (d) between Zap70 tSH2 and LIC-CD3z-YFP channel. Each trace is one cell. Scale bar = 3 µm.

In summary, the reconstitutions experiments demonstrated that the phosphorylation of CD3ζ ITAMs only required functional Lck and that CD3ζ phosphorylation was increased upon LIC-CD3z clustering, potentially because Lck could transiently interact with CD3ζ via its SH2 domains.

### Diffusion analysis of Lck

Because we could not detect clear Lck colocalization with LIC-CD3z clusters, we used diffusion measurements to map Lck interaction with monomeric and clustered CD3ζ and reasoned that any intermolecular interaction would result in slower diffusion of Lck. To avoid emission crosstalk from GFP to mcherry channel as shown in **Supplementary Fig. 2a**, we performed single-point fluorescence spectral correlation spectroscopy as described previously (Benda et al., 2014). The resulted showed that the diffusion times of Lck GFP and clustered LIC-CD3z were on millisecond and sub-seconds time scales for diffraction limited confocal spot. Cross-correlation analysis did not show any significant cross-correlation (**Supplementary Fig. 3a**). However, due to the low signal to noise ratio of the data and cell heterogeneity this finding does not exclude that small (less than 10%) populations do interact. Because the single-point FCS cannot follow the expected spatial heterogeneity within the plasma membrane, we used Line-scanning fluorescence spectral correlation spectroscopy (Line-scanning FSCS) (Benda et al., 2015, Benda et al., 2014) that allowed us to sample a larger area of the plasma membrane compared to single-point FCS with sufficient temporal resolution and self-calibrating properties. The overall line averaged spatial-temporal correlation data show that the diffusion of Lck cannot be fitted by a single component free 2D diffusion model (**Supplementary Fig. 3b, c**). We have thus extracted the average diffusion time of Lck as the time at which the amplitude of the correlation curve drops to its half (**Fig. 7a** and **Supplementary Fig. 3d**). The time evolution of line profiles suggested that the LIC-CD3z clusters are stable over the course of the Line-scanning FSCS measurement. This allows to investigate whether the diffusion of Lck outside or inside the clusters varies. Each selected line scan was split into 20 segments and for each segment we obtained the diffusion time of Lck-GFP and intensity of mCherry channel, which is indicative to the location of LIC-CD3z clusters. The analysis showed that the diffusion coefficient of Lck was highly heterogeneous along the scanned line (black line of **Fig. 7b**). The scatter plot of the intensity distribution of LIC-CD3z and Lck diffusion coefficients along the line showed no apparent correlation (**Fig. 7c**), suggesting that Lck diffusion was not impacted by the presence of LIC-CD3z clusters. Similar results were obtained for signaling-incompetent LIC-CD3z Y-L and monomeric LIC-CD3z-delCry2. This result implied that Lck did not form stable interactions with LIC-CD3z clusters, indicating that phosphorylation most likely arose from transient interactions.

**Figure 7.**
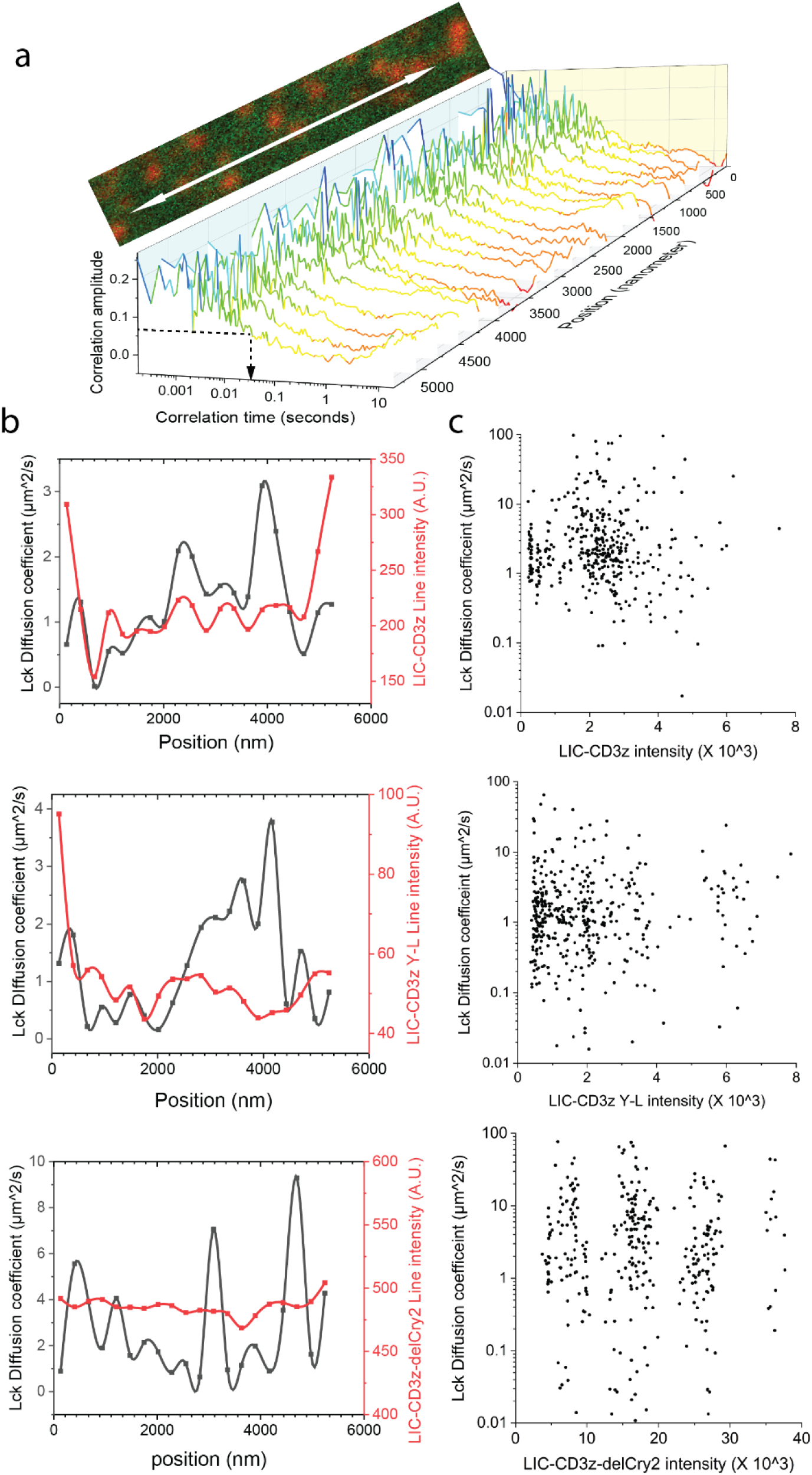
Diffusion analysis of Lck in the absence or presence of LIC-CD3z clusters by Line-scanning FSCS in COS-7 cells. CD3 signaling was reconstituted in COS-7 cells by co-expressing LIC-CD3z and wild-type Lck GFP. **a**. Line-scanning FSCS was performed across the basolateral plasma membrane for 5.4 µm by 488 nm and 594 nm excitation as a bidirectionally scan at 1.3 kHz. A region of interest underneath the cell nucleus that contains LIC-CD3z clusters (red) and Lck GFP (green) as illustrated on the cropped image was chosen to perform the Line-scanning FSCS. The scanned line of the Lck channel was segmented into 20 equal parts for which the autocorrelation curves were plotted. Color indicates correlation amplitude, from which the lag time at half the amplitude of *G(0)* was used to estimate Lck diffusion coefficient as indicated by black dotted arrow in the plot. **b**, Representative line intensity profile (red line) of LIC-CD3z (top), LIC-CD3z Y-L(middle) or LIC-CD3z-delCry2 (bottom) and estimated diffusion coefficients of co-expressed Lck GFP (black line) along the scanning line. n=20 cells. **c**. Scatter plot of 400 intensity values of LIC-CD3z (top), LIC-CD3z Y-L(middle) or LIC-CD3z-delCry2 (bottom) versus estimated Lck diffusion coefficients of the same positions along the line.

## Discussion

We designed an optogenetic tool to test how CD3ζ clustering *per se* impacts on signaling. LIC-CD3z did not trigger signaling via endogenous components of the TCR-CD3 complex since the transmembrane domain of CD3ζ that is known to interaction with other subunit (Call et al., 2002) was replaced with the membrane anchor of Lck. Further, LIC-CD3z is likely to be monomeric when expressed in the plasma membrane because the formation of CD3 ζ-ζ dimer is known to be coordinated through hydrogen bonding of residues on the transmembrane domains of ζ chain (Call et al., 2006). In addition, single molecule studies have shown that the diffusion of Lck10 membrane-anchored proteins fit a free diffusion model (Douglass and Vale, 2005, Lommerse et al., 2006), and co-expression of Lck membrane-anchored GFP and mCherry displayed no colocalization as indicated by a lack of Förster resonance energy transfer (FRET) (Triffo et al., 2012). Light-induced clustering of LIC-CD3z in Jurkat cells showed that the monomer-to-cluster transition of the intracellular domain of CD3ζ was sufficient to trigger signaling events. This was particularly obvious when comparing the effect of light-treated LIC-CD3z to light-treated LIC-CD3z-delCry2 that did not cluster. In reconstituted non-hematopoietic cells, we showed that Lck can phosphorylate monomeric CD3ζ. When basal Lck activity was kept in check, the level of phosphorylation was further increased upon CD3ζ clustering. In T cells, where the basal level of phosphorylation is naturally suppressed by phosphatase activity, clustering of CD3ζ was sufficient to shift the kinase-phosphatase balance and induce signaling that resembles TCR triggering.

The results of the current study do not neatly fit the conformational change model of T-cell activation (Lee et al., 2015, Ma and Finkel, 2010). Such models require tight and rigid association of the TCR αβ-chains to the CD3 signaling chains. For example, it has been proposed that the force of ligand receptor binding is transmitted as a torque from the extracellular to the intracellular signaling motifs (Kim et al., 2012). LIC-CD3z lacks both extracellular and transmembrane domains, and the signaling was initialized from cross-linking of the intracellular domain of CD3ζ chains in a ligand-free manner. Given that the Cry2 domain was linked to the distal C-terminal end of the intrinsically disordered CD3ζ chain (Sigalov et al., 2004) with a flexible linker, we do not expect that clustering of Cry2 generated substantial pulling force to alter the juxtamembrane conformation of CD3ζ chains as was proposed in support of the mechanical force switch model (Lee et al., 2015). It is possible that the lateral clustering of CD3ζ chains caused the cytosolic tails to disassociate from the plasma membrane as proposed by the safety on model (Xu et al., 2008). Such dissociation is more likely to occur *via* molecular crowding within clusters than mechanical forces induced by clustering. However, a previous *in vitro* study has showed that clustering of CD3ζ did not change its random-coil-like unstructured status (Sigalov et al., 2004). It is likely that LIC-CD3z remains dynamically membrane associated (Zimmermann et al., 2017) regardless of its clustering state. It should be also considered that previous work showed that ITAMs can still be phosphorylated by Lck in membrane-bound state (Zhang et al., 2011). Further, a fully phosphorylated CD3ζ chain can remain membrane-associated due to the clusters of positively charge motifs on the ζ-chains (Sigalov et al., 2006).

The mechanism of TCR triggering observed by LIC-CD3z provides hints to the activation mechanism of the TCR-CD3 complex. The idea that TCR-CD3 complexes can be activated by oligomerization directly stems from the observation that oligomeric ligands are required to trigger TCR activation (Boniface et al., 1998, Cochran et al., 2000, Reich et al., 1997). It is also well documented that antibody cross-linking of single chain chimeric receptor of ζ or ε chain of TCR, or γ chain of Fc receptor alone was sufficient to trigger T cell activation (Irving and Weiss, 1991, Romeo and Seed, 1991, Letourneur and Klausner, 1992). Increased ITAM phosphorylation upon CD3ζ clustering has also been reported for *in vitro* systems containing Lck and CD45 (Furlan et al., 2014, Hui and Vale, 2014). In these experiments, CD3ζ phosphorylation may have been driven by the exclusion of CD45 from clustered CD3ζ. Here, we showed that clustered CD3ζ is a more efficient substrate for Lck than monomer CD3ζ if basal Lck activity is kept low. Thus, the enhanced efficiency of Lck and exclusion of CD45 in TCR clusters could work synergistically in T cells. Basal ITAM phosphorylation levels and Lck activity are controlled in T cells by phosphatases such as CD45 (Davis and van der Merwe, 2006) and the kinase Csk (Chow et al., 1993), respectively. The corollary of low basal activity is that substrate clustering can then enhance Lck phosphorylation efficiency independently from CD45 levels. This may be a mechanism of how activation thresholds are set relative to basal states.

Currently, it is not known why Lck phosphorylation of ITAM domains occurs more efficiently in CD3ζ clusters. We found no evidence of stable interactions between Lck and clustered LIC-CD3z. The time Lck spent in the clustered regions was comparable to that spent in non-clustered regions of LIC-CD3z, suggesting that Lck diffusion was not slowed down by LIC-CD3z clusters. It remains possible that Lck interacts more frequently or for longer with clustered LIC-CD3z *via* its SH2 domain but in our experiments, these changes were not detectable.

In summary, we showed that the spatial clustering of the intracellular motif of the CD3ζ chain was sufficient to trigger proximal TCR signaling events that are reminiscent of canonical TCR activation. Lck-mediated phosphorylation of clustered CD3ζ may be a distinct TCR triggering mechanism that is independent of phosphatase activity but requires a low basal level of ITAM phosphorylation. This TCR triggering mechanism is not exclusive of other TCR triggering mechanisms and indeed, may co-exist with other process that induce and regulate TCR signal initiation and propagation.

## Acknowledgement

KG acknowledges funding from the Australian Research Council (CE140100011 and LE180100157) and the National Health and Medical Research Council of Australia (APP1155162 and APP1183588). YM acknowledges the Earlier Career Fellowship funding from the National Health and Medical Research Council of Australia (APP1139003). AB acknowledges ERD Fund-Project No. CZ.02.1.01/0.0/0.0/16_013/0001775. JG acknowledges funding from the National Health and Medical Research Council of Australia (APP1163814). We thank the Biomedical Imaging Facility (BMIF) at the University of New South Wales for providing the instruments and technical support. Especially thank to Dr Michael Carnell for helpful discussions on image Pearson coefficient analysis. We also thank Dr Sophie Pageon for helpful discussion at early stage of the project, Dr Jieqiong Lou for providing the CRISPER-Cas9 LAT knock out Jurkat cell line, and Dr Zhengmin Yang for technical assistance in the laboratory.

## Methods and Materials

### Plasmids

All constructs and mutants used in the current study were prepared by standard subcloning, overlapping PCR and site-directed mutagenesis. The primers used and the sequence of LIC-CD3z are provided in Supplementary tables. The Y to L mutations of the CD3ζ intracellular domain was subcloned from a CD3ζ-6YL-PSCFP2 construct (Pageon et al., 2016). The G-GECO (#32447) and Zap70 tSH2(#27137) constructs were obtained from Addgene.

### Cell culture and light treatment

COS-7 (ATCC CRL-1651) cells were cultured in Dulbecco’s Modified Eagle’s medium (DMEM) with 10% fetal bovine serum (FBS). Jurkat 76 (Pageon et al., 2016), JCam1.6 (ATCC CRL-2063), Jurkat P116 (Williams et al., 1998), Jurkat LAT KO (Gift from JieQiong) cell lines were all cultured in Roswell Park Memorial Institute medium (RPMI) supplemented with 10% FBS and L-Glutamine. Cells were cultured in incubators set at 37 °C with 5% CO_2_. All cell lines were tested and confirmed to be *Mycoplasma* negative. Transient cell transfection was conducted using an electroporator (Invitrogen Neon) following the manufacturer’s protocols and all cells were imaged or treated 18–24 h post transfection. For light activation of LIC-CD3z transfected cells, the cells were washed and light activated in HBSS buffer containing Ca^2+^ and Mg^2+^ (Thermo Fisher, #14025076). For activation on the microscope, the activation was done using a 458 nm or 488 nm laser. For activation of a large cell population for western blot, light activation was done using a 29 mW/cm^2^ 470 nm LED blue light illuminator box (Maestrogen, #LB16) (measured at 488 nm using a Newport photometer).

To measure background phosphorylation levels in COS-7 cells, 25 µM of PP2 (Sigma Aldrich, #p0042) was added to the cell culture media following cell transfection and incubated overnight. The cells were switched to PP2-free HBSS buffer for 1 minute prior to light treatment to allow for the recovery of Lck kinase activity.

### Western blotting

Transfected and wild type Jurkat 76 cells were pelleted by centrifugation at 1500 rpm for 5 min and resuspended in 150 µl of HBSS buffer containing Ca^2+^ and Mg^2+^. The samples were either kept in the dark or light-activated using the LED blue light illuminator box for 45 seconds, followed by further incubation (0 – 5 min) in the dark. All cell treatments here were performed at room temperature (23°C). After incubation, cells were quickly lysed in 8% Lithium dodecyl sulfate LDS loading buffer preheated at 85°C (Invitrogen Cat#NP0007) and incubated with stirring at 500 rpm for 5 min at 85°C (Eppendorf, ThermoMixer C). Lysates were either sonicated (Sonifier 250, Branson) or homogenized with a Bio-Gen PRO200 homogenizer with an attached 5mm X 75mm flat bottom generator probe (Pro-Scientific) and then loaded into 4-12% Bolt Bis-Tris gels (Invitrogen) for gel electrophoresis. Resolved proteins were transferred onto a PVDF membrane (IB24001) using the iBlot2 dry transfer system (Life Technologies).

For Western blotting, membranes were blocked with 5% (w/v) of BSA for 1 hr, followed by overnight incubation in primary antibodies against pERK (1:2000, Cell Signaling, Cat# 9101S), pZAP70 (pY319, 1:1000, Cell Signaling, Cat# 2701S), pCD3ζ (pY142, 1:1000, abcam, Cat# ab68235), pPLCγ (pY783, 1:1000, Cell Signaling, Cat#2821s), GAPDH (1:5000, Abcam, Cat# ab8245) at 4°C in 4% BSA. Blots were then incubated in HRP-linked secondary antibody against mouse or rabbit IgG (1:3000, Cell Signaling, Cat# 7074S and 7076S) in 4% BSA for 1 hr at room temperature. Protein bands were visualized by chemiluminescence (Pierce ECL Western Blotting Substrate, Thermo Scientific) following the manufacturer’s protocol and then imaged using ImageQuant LAS4000 Mini gel documenter (GE Life Sciences). Protein band intensity was analysed in ImageJ (National Institutes of Health) by taking the integral of the pixel intensity of the protein bands.

### Imaging

Cells were imaged in either a Leica SP5 or Zeiss 880 laser scanning Confocal microscope. For multicolour imaging, sequential scanning mode was used to avoid crosstalk during the collection of emitted fluorescence. For Ca^2+^ imaging of Jurkat cells, transfected cells were washed and imaged at room temperature in HBSS buffer containing Ca^2+^ and Mg^2+^ through an HCX APO L 20X/1.0 NA water immersion objective. G-GECO and LIC-CD3z were sequentially excited by 458 nm and 594 nm lasers, and fluorescence emission was collected between 470 – 540 nm and 590 – 670 nm spectral bands, respectively. For Zap70 COS-7 live cell experiments, transfected cells were washed and imaged in HBSS buffer containing Ca^2+^ and Mg^2+^ using the Zeiss 880 with an LD C-Apochromat 40x/1.1 NA water immersion objective at room temperature. GFP, YFP and mCherry that tagged to Lck, LIC-CD3z-YFP and Zap70 tSH2, respectively were excited by 458 nm, 514 nm and 594 nm lasers, which were sequentially reflected to the objective through the triple dichroic mirror (458/514/594 nm). Two PMTs and one GaAsP detector were gated to 460 – 508 nm, 526 – 580 nm, and 605 – 721 nm to sequentially capture the corresponding fluorescence. For pCD3z immunostaining, COS-7 cells co-transfected with LIC-CD3z or LIC-CD3z-delCry2 with Lck GFP were light treated/not treated and fixed in 4% paraformaldehyde and permeabilized with 0.05% Triton X-100. The cells were blocked with 4% BSA and stained with a 1:200 (v/v) dilution of Alexa Fluor 647 Mouse anti CD247 pY142 antibody (BD Bioscience #558489) for 1 hour at room temperature. The stained cells were washed and imaged on the Leica SP5 with sequential excitation by a 488 nm, 561 nm and 633 nm pulsed white light laser directed at an HCX PL APO CS 20x 0.70 IMM objective through of 488/561 nm dichroic mirror and 633 nm notch filter. Two hybrid HyD and one PMT detectors were spectrally gated to 497 – 543 nm, 569 – 630 nm, and 652 – 774 nm to sequentially capture the corresponding fluorescence. A line average of 32 lines was used during image aquisition to increase the signal to noise ratio of the image. The two colour imaging of LIC-CD3z and Lck GFP in COS-7 cells were acquired under Airyscan mode on Zeiss880 microscope. The tagged GFP and mCherry fluorophores are excited frame by frame sequentially using 458 nm and 594 nm lasers that reflected from the triple dichroic mirror (458/514/594 nm) to the Plan-Apochromat 63x/1.4 Oil DIC M27objective. The emission collected travel through the same dichroic mirror and a dual emission band pass filter (bandpass 495-550 nm/ LP 570) prior arrive to the Airyscan fibre head, which were set to super-resolution mode under Zen software. The diameter of the hexagon was configured to be equivalent of 1.26 Airy unit. The super-resolution image was reconstructed using Zen software prior to colocalization analysis using self-written MATLAB code.

### Image analysis

Majority of the image analysis were conducted in custom wrote MATLAB code (Mathworks). The script is available online at (https://github.com/mayuanqing8/Light-induced-CD3-clustering-project).

The Ca^2+^ profile of individual cells shown in **Fig. 2c** were extracted from the time-lapse images by manually selecting 20 regions of interest (ROI) and plotted as a function of time in ImageJ. The increase of Ca^2+^ shown in **Fig. 2d** was quantified as the normalized increase of G-GECO intensity (average of the last 6 frames subtracted by the average of the first 6 frames divided by the average of the first 6 frames).

The quantification of pCD3AF647 immunostaining shown in **Fig. 4** was performed as follows: Since not all the cells were co-transfected with LIC-CD3z and Lck GFP, the first step was to identify the cells containing both constructs. This was done by logical AND selection of pixels in both LIC-CD3z and Lck GFP channels that had intensity values greater than the ostu threshold value of the respective channel. This produced a mask to select cells that contained both constructs. The same mask was used to extract the intensity values of the pCD3AF647 channel. In order to correct for the intensity variation of pCD3AF647 due to difference of protein expression level between LIC-CD3z and LIC-CD3z-delCry2, the extracted mean intensity value of pCD3AF647 was divided by the mean intensity value of LIC-CD3z or LIC-CD3z-delCry2. In order to correct for intensity variation due to human error, such as change of objective focus, or cell staining efficiency, the background intensity of pCD3AF647 from cells that contained only LIC-CD3z with no Lck GFP was used to standardize pCD3AF647 intensity across different sample groups.

The cytosol to plasma membrane translocation of Zap70 tSH2 shown in **Fig. 5c** is analysed as follows: briefly, ROIs containing single cells were first manually cropped from the time-lapse movies using ImageJ and saved as hyperstack Tiff images, which are load into MATLAB. The borders of each cell were identified by Sobel edge detection, followed by a number of clean-up steps including image dilation, filling interior holes, and smoothing of the detected object. An ascending (0-1) distance to centre 2D matrix was created from the centre of mass to identified edge of cells. Masks for cell plasma membrane and cytosol was created by logical selection of matrix values between 0 to 0.25 and 0.25 to 1, respectively. The fluorescence intensity of the Zap70 tSH2 in the selected regions was extracted and analyzed. To verify the code works, the same mask was used to extract intensity change of permanently membrane attached LIC-CD3z, which has showed no change as shown in **Fig. 5c** and **Supplementary Fig. 1**.

For image Pearson coefficient analysis shown in **Fig. 6b, d**, the images were first thresholded using otsu (Pike et al., 2017) method to remove background regions as Pearson coefficient analysis method is known to be sensitive to background noise (Costes et al., 2004). The image Pearson coefficient was calculated as 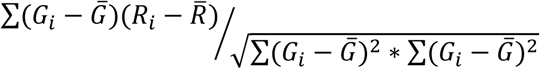, where *G*_*i*_, *R*_*i*_ and 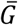 and 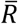 are incident and mean pixel intensity values of green and red channels of the background removed regions.

### Fluorescence Spectral correlation Spectroscopy

Both the single point FSCS and Line-scanning FSCS was acquired using the Zeiss 880 laser scanning confocal microscope. 488 nm and 594 nm CW lasers were focused on the Plan-Apochromat 63x 1.4 oil objective through a dual colour dichroic mirror (488/594 nm) to simultaneously excite GFP and mCherry at the basolateral membrane of the cell. The closer distance of detection volume from the coverslip helps to reduce optical distortion related to refractive index mismatch, affecting the correlation curve (Levitus, 2011). The fluorescence emitted was collected by the same objective and dichroic mirror prior arrive to the GaAsP detector set at either 6 spectral channels (495-510nm, 511-530nm, 531-560nm, 561-610nm, 611-630nm, 631-680nm) for single-point FSCS or 10 spectral channels (481-499nm, 500-518 nm, 519-538 nm, 539-557 nm, 558-576 nm, 577-595 nm, 596-616 nm, 617-635 nm, 636-654 nm, 655-673nm) in single photon counting detection mode. For Line-scanning FSCS, the scanning distance was 5.355 micrometers over 256 pixels with a pixel size of 21 nm. The laser was scanned in a bidirectional manner over the same one-dimensional space at a speed of 0.51 μs per pixel, and 154 μs line scan time. 300,000 lines were collected over a time period of 47 seconds. The data was saved and converted into TTSR file format in a custom made LabView software (Benda et al., 2014). The analysis of single point FSCS data was performed as previously published (Benda et al., 2014), and the Line-scanning FSCS data was analysed similar to single point FSCS analysis with some modifications. Briefly, the pixel and spectral channel identity of each photon collected is used to retrieve where the photon was detected from. The assumption was that in any given time, the intensity of any spectral channel was attributed from a linear combination of the fluorescent species present in the excitation volume. Based from the emission spectra of each reference dye (the spectra referred is the dichroic mirror filtered spectra) and the dye mixed sample spectra, one can estimate the contribution from respective dye species by applying a spectral filter *f*_*gi*_ and *f*_*ri*_ to the sample spectra, *f*_*gi*_ and *f*_*ri*_ are the calculated filter for GFP (*g* stand for green) and mCherry (*r* stand for red), respectively. Essentially, the multiplication of the sample spectra with the calculate filter should reproduce the reference dye spectra. Here the spectra are the intensity normalized spectra, and the sum of the two filters is equal to 1. The filters are recalculated for each sample and treated as a constant value in the correlation analysis. The calculated filters are integrated into the autocorrelation algorithm to weight the photons depend on the spectral channel to recover the correlation that attributed from individual dyes. The cross-correlation is calculated as: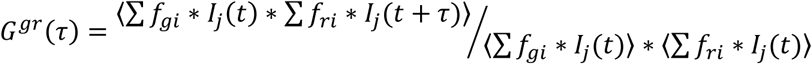. Here ⟨ ⟩ notes averaging over all time *t. f*_*gi*_ and *f*_*ri*_ are calculated filters applied to green and red channel respectively. *I*_*j*_*(t)* is the photon intensity of spectral channel *j* at time *t*. In case of autocorrelation, filter *f*_*gi*_ *= f*_*ri*_. Compared to traditional single-point FCS, the sampling rate of Line-scanning FSCS is slower and defined by the pixel dwell time of 0.51 μs, and the probability of multiple photon event during this time period is considerable. Under single photon counting mode, single photon events are registered as a pixel intensity of 1, where multiple photon event re registered 2, 3… and so on, from 0 to 255 as 8-bit Tiff images. Higher weighting factors (scaling the filter by the intensity value) has to be applied to those multiphoton pixels so that their contribution to the correlation calculation are scaled accordingly.

The fitting of autocorrelation curves was performed as previously described (Benda et al., 2015). Here, the autocorrelation took a two-dimensional spatial and temporal correlation form, where the Gaussian shaped lateral spread in the scanning axis was calculated by the spatial lateral cross correlation of the neighbouring pixels along the line at *G(0)*. The Gaussian here is equivalent to the cross section of the product of the excitation and detection PSF of the microscope (Benda et al., 2015). The FWHM of the Gaussian curve from the spatial cross-correlation was used to conveniently retrieve the diameter *w* of the detection volume, so that no calibration using a reference dye is needed. Orthogonal to the plane of the Gaussian form was the temporal autocorrelation curve that can be analyzed with model fitting or fitting free options. The intensity fluctuation of 256 pixels along the line can be split into 20 segments each of a size of 250 nm that equivalent of 20 raw single-point FCS measurements. For model fitting of the correlation curves, the autocorrelation curves of the 20 segments are averaged to increase statistical robustness. Here, the resulting 2D autocorrelation curve can be fitted to the two-dimensional n species free diffusion model as shown in **Supplementary Fig. 3b, c**. 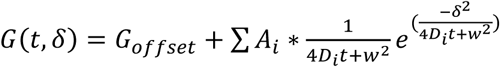, where *t* is the temporal correlation lag time, and *δ* is spatial correlation lag. *A*_*i*_ and *D*_*i*_ are the autocorrelation amplitude and diffusion coefficients of the respective species. *w* is the beam waist of excitation laser.

For investigating the heterogeneity of Lck GFP diffusion along the scanning line, the 20 raw auto-correlation curves along the line were analyzed individually. Given that a single species diffusion model was insufficient to fit Lck GFP autocorrelation curves as shown in **Supplementary Fig. 3b, c**, and the signal to noise ratio of the segmented single-point autocorrelation curves are too low for accurate two species model fit. We used a fitting free method as shown in **Supplementary Fig. 3d and Fig. 7a**, where data of each segment was subject to independent temporal correlation analysis. Cubic spline smoothing was applied to each correlation curve to extrapolate to zero lag time to estimate *G*(0). The *G*(0) value was used as a reference to interpolate the half lag time, who’s amplitude is 50% of the value of *G*(0). This is defined as the transition time *γD* of the molecule across the excitation volume. Here Lck GFP diffusion coefficient is calculated as 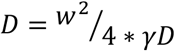. The beam waist *w* value was obtained from the spatial correlation analysis as mentioned above. In this fitting free analysis, there was no assumption made relating to the number of diffusion component or diffusion mode.

