## Supplementary table, figures and movie legend for "Clustering of CD3ζ is sufficient to initiate T cell receptor signaling"

**Supplementary tables:**

**Protein sequence of LIC-CD3z and mutants:**

MGCVCSSNPE_GGGGD_RVKFSRSAEPPAYQQGQNQL**Y**NELNLGRREE**Y**DVLDKRRGRDPEMGGKPRRKNPQEGL**Y**N
ELQKDKMAEA**Y**SEIGMKGERRRGKGHDGL**Y**QGLSTATKDT**Y**DALHMQALPPR_GSGGGGSGT_MVSKGEEDNMAIIK
EFMRFKVHMEGSVNGHEFEIEGEGEGRPYEGTQTAKLKVTKGGPLPFAWDILSPQFMYGSKAYVKHPADIPDYLKLSFP
EGFKWERVMNFEDGGVVTVTQDSSLQDGEFIYKVKLRGTNFPSDGPVMQKKTMGWEASSERMYPEDGALKGEIKQRL
KLKDGGHYDAEVKTTYKAKKPVQLPGAYNVNIKLDITSHNEDYTIVEQYERAEGRHSTGGMDELYK_GSTGSTGT_MK
MDKKTIVWFRRDLRIEDNPALAAAAHEGSVFPVFIWCPEEEGQFYPGRASRWWMKQSLAHLSQSLKALGSDLTLIKTH
NTISAILDCIRVTGATKVVFNHLYDPVSLVRDHTVKEKLVERGISVQSYNGDLLYEPWEIYCEKGKPFTSFNSYWKKCL
DMSIESVMLPPPWRLMPITAAAEAIWACSIEELGLENEAEKPSNALLTRAWSPGWSNADKLLNEFIEKQLIDYAKNSKK
VVGNSTSLLSPYLHFGEISVRHVFQCARMKQIIWARDKNSEGEESADLFLRGIGLREYSRYICFNFPFTHEQSLLSHLRFFP
WDADVDKFKAWRQGRTGYPLVDAGMRELWATGWMHNRIRVIVSSFAVKFLLLPWKWGMKYFWDTLLDADLECDIL
GWQYISGSIPDGHELDRLDNPALQGAKYDPEGEYIRQWLPELARLPTEWIHHPWDAPLTVLKASGVELGTNYAKPIVDI
DTARELLAKAISRTREAQIMIGAAPDEIVADSFEALGANTIKEPGLCPSVSSNDQQVPSAVRYNGSKRVKPEEEEERDMK
KSRGFDERELFSTAESSSSSSVFFVSQSCSLASEGKNLEGIQDSSDQITTSLGKNGCK*

The key elements of the construct are underlined in order as: Lck10, CD3ζ intracellular domain, mCherry, photolyase homology region (PHR) 1-498 AA of the Cry2. Linker sequences are shown between key elements. Tyrosine to Leucine mutations in LIC-CD3z Y-L construct are shown in bold red. In the construct LIC-CD3z-YFP, mCherry was replaced by YFP (monomeric Venus). In LIC-CD3z-delCry2, the Cry2 sequence was replaced by a stop codon.

**Primers list:**

| Primer name | Primer sequence (5’ to 3’) |
|---|---|
| Lck10 OVLP RV | atcccctcctcctccttcagggtttgagctgcagacacatcccatggtggcctcgagatctgagtc |
| Zeta cyto OVLP FW | aggaggaggaggggatagagtgaagttcagcaggagcgcaga |
| Zeta cyto KpnI RV | atggtaccacttccaccgcctccagaaccgcgagggggcagggcctgcatg |
| mCherry KpnI FW | agtggtaccatggtgagcaagggcgaggagga |
| mCherry BsrGI RV | cttgtacagctcgtccatgccgccggtggagt |
| Cy2PHR BsrGI FW | gctgtacaagggctcaactggaagtacaggaacaatgaagatggacaaaaagacta |
| Cy2PHR NotI RV | gtcgcggccgctcatttgcaaccatttttccca |
| mCherry stop NotI RV | gtcgcggccgctcagtacagctcgtccatgccgccggtggagtgg |
| Venus KpnI FW | agtggtaccatggtgagcaagggcgaggagc |
| Venus BsrGI RV | ccttgtacagctcgtccatgccggcggcggtcacgaactccagca |
| Lck R154K RV | tctcgctctccttgatgaggaaggagccgtga |
| Lck R154K FW | tccttcctcatcaaggagagcgagagcaccgc |
| Lck K273R RV | cttcaggctccgcaccgccaccttcgtgt |
| Lck K273R FW | gtggcggtgcggagcctgaagcagggca |

**Supplementary figures:**

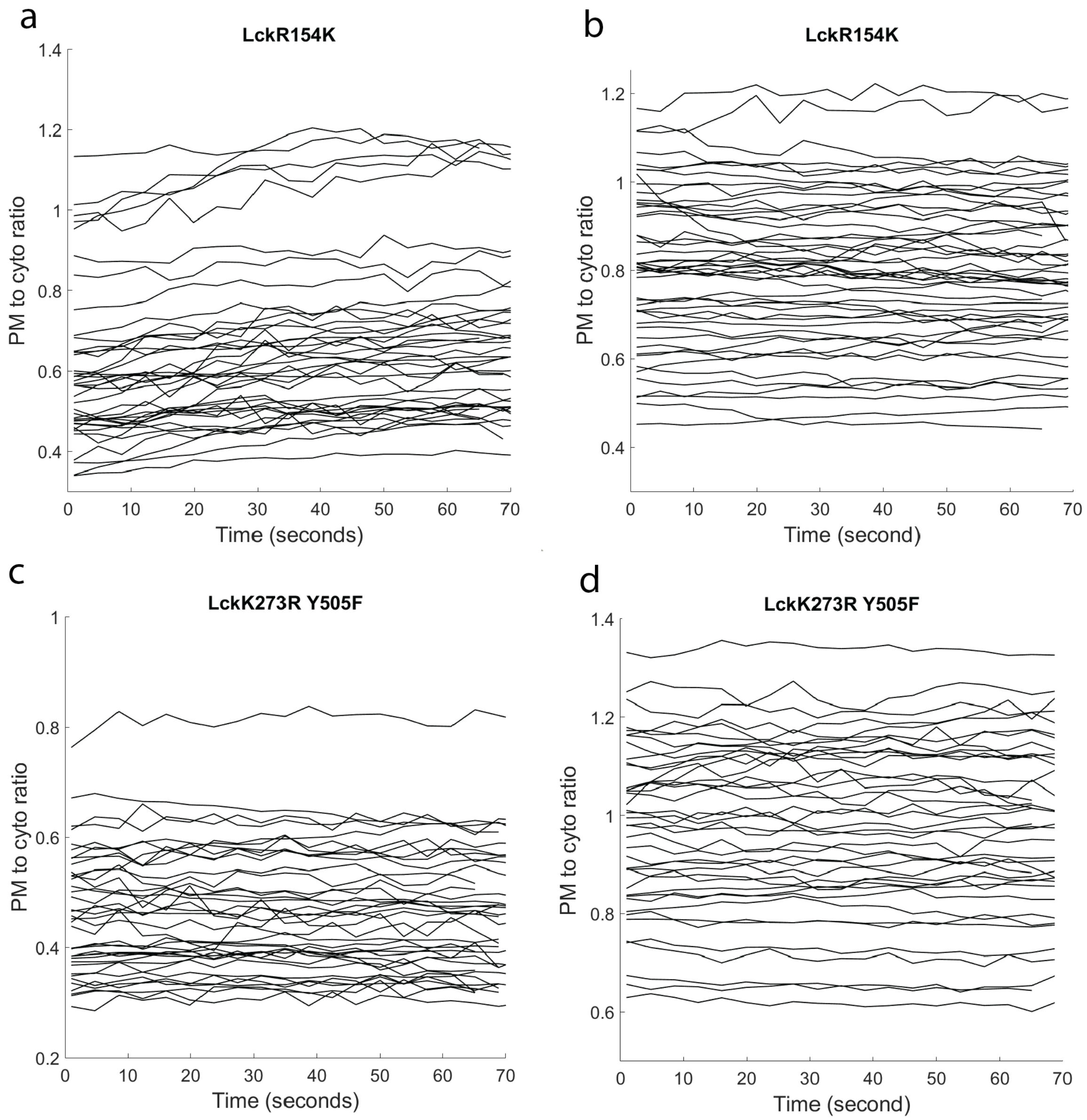


**Supplementary figure 1**. Plot of the cell plasma membrane to cytosol distribution ratio of Zap70 tSH2 (left) and LIC-CD3z-YFP (right) over time after light induced clustering of LIC-CD3z-YFP in cells co-transfected with LIC-CD3z-YFP, Zap70 tSH2 and the indicated Lck mutant, including SH2 domain binding mutant (Lck R154K) (a, b) and kinase dead mutant (Lck K273R Y505F) (c, d). n=40, each line represents one cell.

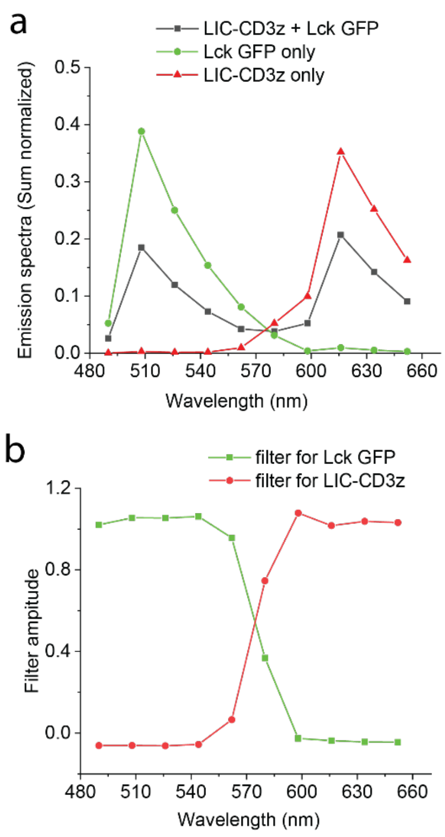


**Supplementary figure 2. Calculated spectral filter for Lck GFP and LIC-CD3z in Line-scanning FSCS a**, Emission spectra of Lck GFP (green) and LIC-CD3z (red) in transfected COS-7 cells were excited simultaneously by 488 nm and 594 nm lasers, which resulted in a mixed sample spectra (black). The fluorescent signal was collected by the GaAsP detector on the Zeiss 880 Laser scanning confocal microscope set at 10 channel mode with 17.8 nm per channel width. **b**, the calculated spectral filter for Lck GFP (green) and LIC-CD3z (red) was used to extract photons of respective species in the correlation analysis.

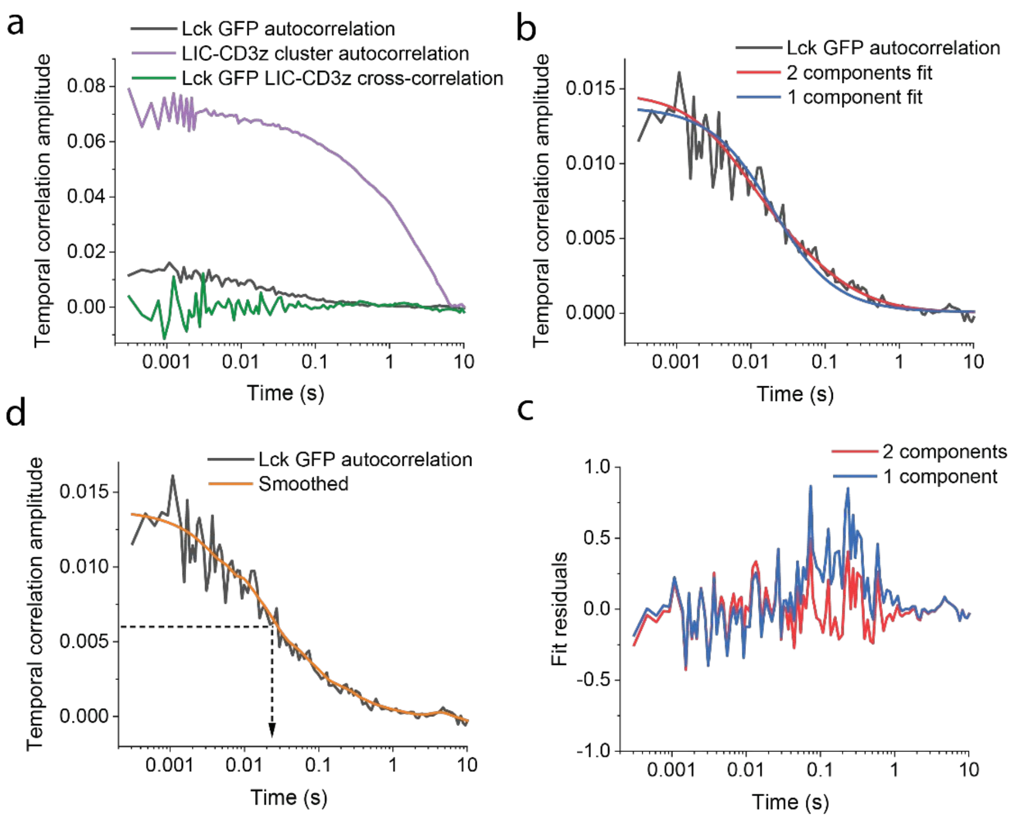


**Supplementary figure 3**, **a**, Representative autocorrelations and cross-correlation of Lck GFP and LIC-CD3z clusters as indicated. **b**, representative model fit of the Lck GFP temporal autocorrelation curve (black) to one component (blue) or two component (red) two-dimensional free diffusion models as described in methods. **c**, the residuals of the corresponding fit shown in b. The residuals are normalized by the standard deviation of the 20 auto-correlation curves along the line. The sum of squared residuals for the 1 component and 2 component fit were 6.70 and 2.79, respectively. **d**, the fitting free approach to estimate the diffusion time of Lck GFP, which is the correlation lag time at half the value of the *G(0)* that extrapolated from the smoothed raw autocorrelation curve.

**Supplementary movies**

**Video 1** Representative movie of light induced $Ca^{2+}$ influx of Jurkat 76 cells co-transfected with LIC-CD3z and G-GECO. Channel shown are G-GECO (green), LIC-CD3z (red), and two channels merged. Video were taken on LeicaSP5 confocal microscope using 20X/1.0 NA water immersion objective under simultaneous excitation by 458 nm and 594 nm lasers. Fluorescence emission was collected between 470 – 540 nm and 590 – 670 nm two spectral bands. The unit of time stamp is in seconds. Scale bar = 400 μm.

**Video 2** Representative two single cell movies of Zap70 tSH2 translocation from cell cytosol to plasma membrane upon light induced clustering of LIC-CD3z-YFP in the reconstituted COS-7 cells. Cells are co-transfected with LIC-CD3z-YFP, Lck GFP and Zap70 tSH2 three constructs showing in three channels from left to right. The video was taken on the Zeiss880 confocal laser scanning microscope using 63x 1.4 NA oil objective focusing at the middle of the cell. The unit of time stamp is in seconds. Scale bar = 10 μm.

**Video 3** Representative Zap70 tSH2 translocation from cell cytosol to plasma membrane upon light induced clustering of LIC-CD3z-YFP in a population of reconstituted COS-7 cells. Cells are co-transfected with LIC-CD3z-YFP, Lck GFP and Zap70 tSH2 three constructs showing in three channels from left to right. The video was taken on the Zeiss880 confocal laser scanning microscope using 40x/1.1 NA water immersion objective focusing at the middle of the cell. The unit of time stamp is in seconds. Scale bar = 30 μm.

**Video 4** Representative movie of Zap70 tSH2 translocate into newly formed LIC-CD3z clusters in reconstituted COS-7 cells. Cells are co-transfected with LIC-CD3z-YFP, Lck GFP and Zap70 tSH2 three constructs showing in three channels from left to right. Image taken by 63x 1.4 NA oil objective focusing at the basolateral membrane of the cell. The unit of time stamp is in seconds. Scale bar = 5 μm.
